# Flexible 3D multimodal organoid interface reveals functional subnetworks and coordinated circuit activity in cortical organoids

**DOI:** 10.64898/2026.10.02.756317

**Authors:** Madison N. Wilson, Elizabeth K. Kharitonova, Joseph Yeh, Joann Lee, Shadee Hemaidan, Samuel Essig, Marie Shi, Colt W. Crain, Mehrdad Ramezani, Fengyi Sun, Teng Zhou, Ikbal Surucu, Farzad Mortazavi, Ertugrul Cubukcu, Martin Thunemann, Ella Zeldich, Duygu Kuzum

## Abstract

The balance between excitatory and inhibitory (E/I) neurons is fundamental to cortical function, and its disruption underlies a wide range of neurological disorders, including epilepsy, Alzheimer’s disease and autism spectrum disorder. Cortical organoids offer a promising model to study this balance, as they recapitulate key features of brain development, including the emergence of E/I circuitry. However, no existing technology can simultaneously resolve neuronal identity, spatial organization, and dynamic network activity within intact organoids, limiting precise interrogation of E/I circuit function. Here, we report INFORM, an INtegrated Flexible Organoid Recording Multimodal neural interface that conformally wraps organoids of any size and shape, enabling long-term recordings over 371 days with above 90% of channels detecting spikes. INFORM interface combines four distinct modalities; electrophysiology, two-photon calcium imaging, targeted electrical microstimulation, and chemical intervention to monitor and modulate neural activity across multiple spatial and temporal scales spanning from action potentials from single neurons to synchronized bursts from large-scale populations. Such high-resolution interrogation enables probing of emergence of functional connectivity in fusing assembloids and resolves spatially discrete functional subnetworks that can be selectively recruited and modulated by INFORM. Neurotransmission blockade confirms that stimulus-evoked suppression of neuronal spiking is mediated by GABAergic inputs providing direct functional evidence of the presence of inhibitory signaling in organoids. This work allows single-cell dissection of functional networks in brain organoids, providing a powerful platform for modelling and studying cortical development, neurological diseases, and the E/I imbalances underlying circuit dysfunction.

## Introduction

Cortical organoids are 3D brain models derived from human pluripotent stem cells that recapitulate key aspects of cortical development, such as regional identity, glutamatergic/GABAergic cell types, and neural oscillations, making them powerful systems to study neural networks in vitro^1–4^. Organoids are particularly valuable in the context of new approach methodologies (NAMs) since they bridge the complexity of in vivo brains with the tractability of 2D cultures and potentially reduce the need for animal models^5, 6^. Advances in neural interface technologies are increasingly being pursued to enable precise real-time monitoring of electrical activity and functional connectivity in brain organoids^7–11^. An important yet understudied application of organoids is investigating excitatory-inhibitory (E/I) balance, which is essential for information processing, network stability, and proper brain function^12–15^, while imbalance has been implicated as one of the mechanisms behind disorders such as epilepsy, Rett Syndrome, Tourette’s Syndrome, Fragile X Syndrome, schizophrenia, and Parkinson’s disease^16–22^. Current approaches to characterize E/I circuits are typically post hoc immunostaining or transcriptomic analyses, which require tissue fixation and labeling, thus impeding longitudinal studies. While they provide detailed and, in some cases, spatial assessment of cell types and synaptic components^23, 24^, these methods fail to capture how connectivity, maturation, and spatiotemporal synaptic dynamics give rise to functional network activity over time. Cortical organoids provide an ideal platform for studying functional E/I network dynamics, yet no current technology can simultaneously capture neuronal identity, precise spatial location, and real-time network activity in 3D.

Conventional 2D microelectrode arrays (MEAs) widely used for organoid recordings cause flattening and deformation of 3D architecture and record only a fraction of the entire organoid surface^1, 10, 25, 26^. Recent efforts have focused on developing MEAs capable of conforming to the three-dimensional architecture of organoids, enabling more comprehensive electrophysiological recordings across the surface^7–11^. However, improvements in mechanical flexibility and electrode density alone are insufficient to address the fundamental challenge of simultaneous multimodal recordings necessary to resolve cellular functional identities, precise spatial coordinates, and temporal dynamics crucial for capturing single cell resolution network activity from large neuronal populations.

In this study, we present ’INFORM,’ a flexible multimodal neural interface (**Fig. 1a-b**) that uniquely combines dense electrical recordings and targeted microstimulation with high-resolution two-photon microscopy to enable multimodal interrogation of 3D organoid neuronal networks at single-cell resolution (**Supplementary Video 1**). The interface demonstrates marked improvements over previous technologies (**Supplementary Table 1**) and is distinguished by two capabilities that prior organoid interfaces have not demonstrated. First, its thin, transparent, flexible profile was co-designed with the electrode geometry and PDMS housing so that four distinct modalities can be applied simultaneously to the same intact organoid. While individual capabilities have precedents in the literature, this combination has not previously been realized, and it is necessary for comprehensive characterization of organoid networks because each modality is limited when used alone. Second, INFORM sustains this multimodal access at weekly intervals for more than one year, the longest and most frequent recording strategy to date, establishing a level of long-term stability that previous technologies have left unaddressed. This combination of enabling frequent longitudinal measurements and multimodal readout is what makes it possible to resolve organoid network dynamics over developmentally relevant timescales.

**Figure 1.**
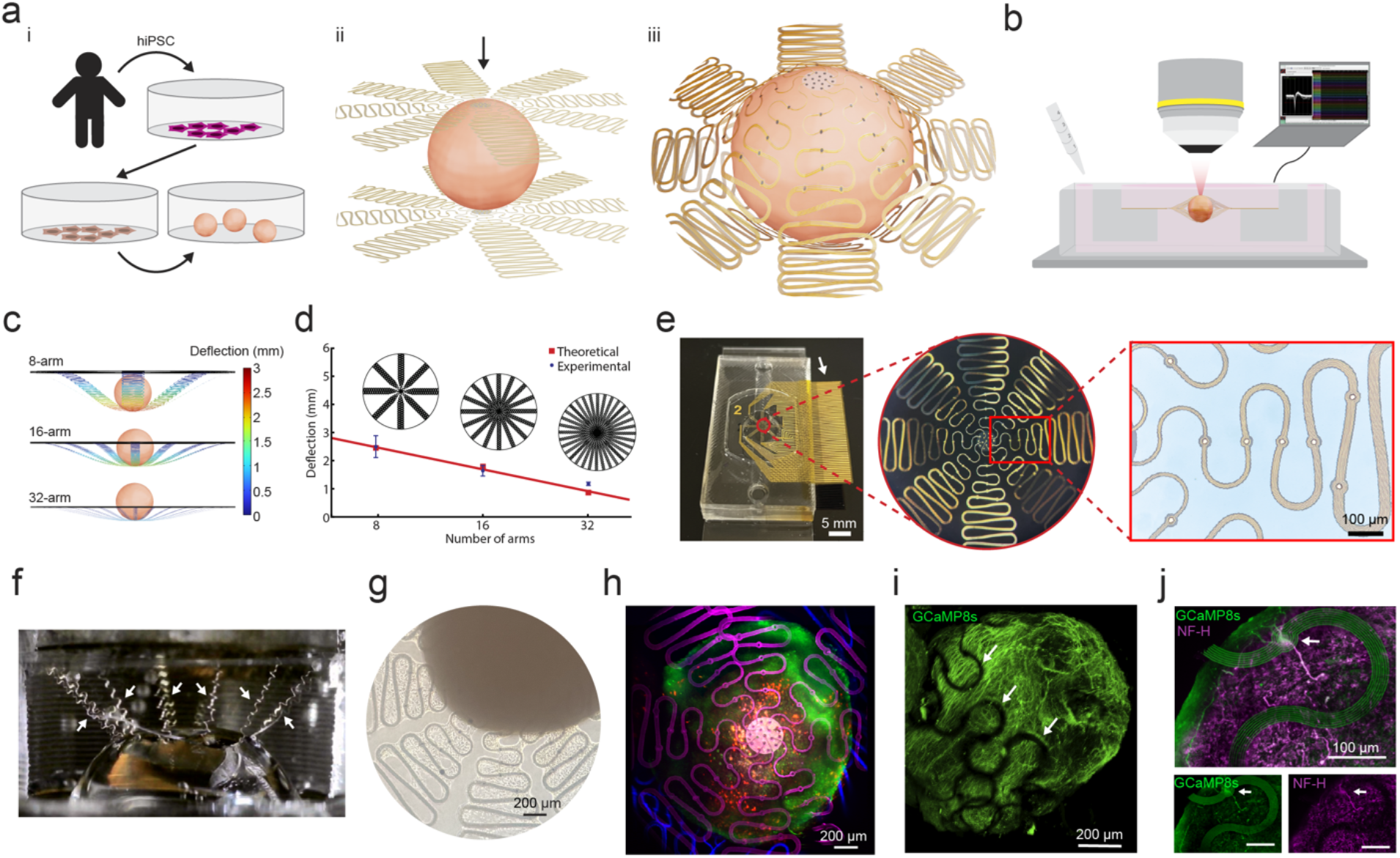
Design and development of the INFORM interface for cortical organoids. (**a**) (i) Generation of human induced pluripotent stem cells (hiPSCs) from skin fibroblasts; differentiation of hiPSCs into neurons which aggregate into cortical organoids. (ii) Organoid is plated on the bottom interface, and another interface is placed on the top (iii) to provide full surface coverage in electrical recordings. (**b**) INFORM interface with the organoids are housed in a PDMS well with ports for media delivery and an open top to permit upright microscopy with simultaneous electrophysiology and microstimulation. (**c**) COMSOL simulations of deflection for different numbers of arms. (**d**) Simulation (red) and empirical (blue) results of deflection distance vs. number of arms. (n = 10 deflection tests conducted per arm condition) (**e**) Photo of an assembled interface in a PDMS well. Callouts show the shape of gold wires at the center of the interface where the organoid will be placed. Arrow points to connection pads which protrude outside the PDMS well to connect via a ZIF connector during recordings. (**f**) Interface deflection by a water droplet. Arrows point to stretched interface arms. (**g**) Neurites are observed to branch along interface arms, showing strong physical integration between the organoid and the interfaces (image taken eleven weeks post-plating). (**h**) Two-photon microscopy image shows z-projection of an organoid between two interfaces. Green is GCaMP8s, red is mCherry labeling neurons, blue is bottom interface and magenta is top interface. (**i**) CLARI-O-cleared organoid showing conformal contact of interface (arrows) to the organoid. (**j**) CLARI-O-cleared organoids immunostained with neurofilament heavy chain (NF-H) shows axons and cells in close proximity to an electrode (arrow).

The interface conforms to curved organoid surfaces of any size and shape, enabling uninterrupted media exchange and undisturbed growth and maturation in long-term experiments. Tight integration with 128 microelectrodes permits detection of spikes from above 90% of the channels while capturing the full spectrum of functional neuronal activity from individual action potentials to large-scale synchronized bursts across organoids and fusing assembloids. We specifically integrated two-photon microscopy imaging with the INFORM interface since its high spatial resolution, deep tissue penetration, and reduced photobleaching complemented the INFORM’s capabilities for long-term visualization of population activity from whole organoids with cellular spatial resolution (**Supplementary Videos 2-4**). We leveraged the high spatiotemporal resolution multimodal recordings and targeted microstimulation capabilities of INFORM to selectively switch neuronal populations on and off, mechanistically dissecting circuit-level activity in ways not possible with existing technologies.

To this end, here, we demonstrated controlled enhancement and suppression of activity in neuronal populations in organoids using the INFORM interface, recapitulating in vivo circuit motifs (**Supplementary Video 5**). Pairing microstimulation with pharmacological GABA-A receptor inhibition and calcium imaging enabled dissection of distinct inhibitory and excitatory circuit-level contributions to organoid network dynamics, beyond the reach of passive or unimodal recordings. Overall, our flexible 3D interface overcomes current technological limitations by enabling longitudinal, multimodal, spatially resolved, functional circuit-level recording and active network modulation within intact human organoids, positioning it as a powerful platform for studying cortical development and neurological diseases.

## Results

### Design, fabrication, and characterization of 3D INFORM interface

Developing interfaces that maximize electrode density within a minimal footprint is essential for enabling concurrent imaging, preserving healthy organoid growth, and achieving longitudinal multimodal recordings. We designed 3D interfaces that conformally wrap the curved organoid surface from the top and bottom through flexible serpentine arms, which permit necessary oxygen and nutrient diffusion while keeping organoids suspended in media (**Fig. 1a**). We generated dorsal cortical organoids from human induced pluripotent stem cells (hiPSC) following a previously established protocol, giving rise to diverse populations of neuronal and glia cells^27^ (**Fig. 1ai, Supplementary Fig. 1, Methods**).

At the age of 5-7 months, the organoids were plated between two INFORM interfaces to allow for attachment, continuous maintenance and weekly recordings for one year without needing to remove or replace organoids between recordings (**Fig. 1aii**). Each interface consisted of 64 electrode channels (128 channels total) placed on the top and bottom organoid hemispheres for high-density recordings across the organoid surface (**Fig. 1aiii**). The channels consisted of 12 μm-diameter electrodes connected to 2-μm-wide gold wires encapsulated in transparent polymer parylene C (**Supplementary Fig. 2**). Narrow traces of conductive lines and small size of electrodes allowed imaging of organoids with minimal interference to cell visibility and calcium transients at different depths from the surface (**Supplementary Fig. 3**). Ultrasmall electrode diameter (12 μm) was chosen to improve spatial selectivity in electrical recordings and microstimulation. The interfaces were housed in a PDMS well with an open top for microscopy, access on the side to electrode connection pads for electrophysiology, and fluidic ports for media delivery during long-term co-culturing of organoids with the interfaces (**Fig. 1b**). Using finite element method (FEM) modeling and experiments, we systematically investigated the mechanical flexibility of serpentine designs across varying arm numbers, widths, and thicknesses. We found that lower number of arms increased deflection under organoid weight (**Fig. 1c**), with experimental measurements using weight-matched organoid surrogates in great agreement with simulations (**Fig. 1d**). Modeling also showed that decreasing interface thickness and arm width increased flexibility and deflection distance (**Supplementary Fig. 4**). Based on simulation results, we chose to design the interfaces with 8 arms (each arm housing 8 metal electrode wires), 4 μm thickness, and a 2 μm margin (distance of outermost electrode wire to the edge of the arm) (**Fig. 1e, Supplementary Fig. 5**). These design parameters yielded highly flexible interfaces (**Fig. 1f**) which could achieve conformal contact with organoids, ensuring reliable electrode-tissue coupling without mechanical failure.

To capture spikes with high fidelity, 12 μm-diameter electrodes were patterned in the center of the interface distributed over a 1.6-mm-diameter area to match the average diameter of organoids plated in this study (**Fig. 1e, Supplementary Fig. 5**). The total PDMS well footprint was small and portable, thus permitting weekly multimodal recordings of electrophysiology on a heated microscope stage outside the incubator (**Fig. 1e**). After assembly of the interfaces with PDMS wells, organoids were plated and left to attach and integrate with the interface for two weeks before recordings began, at which point neuronal projections were seen branching along interface arms, suggesting robust physical integration of the organoid with the interfaces (n = 10 organoids, **Fig. 1g, Supplementary Fig. 6**). That was followed by placement of another interface of the same design over the top of organoids to allow electrophysiological recordings from the upper hemisphere as well (**Fig. 1h**). Organoids and interfaces were positioned to maximize electrode contact with organoid surfaces. Any electrodes not initially covered later formed contact with neuronal projections and signals originating from these electrodes were included in analysis since functional connection was presumed to exist between organoids and their projections. Both two-photon microscopy (**Fig. 1h and Supplementary Fig. 3b**) results and immunostaining on CLARI-O^28^-cleared organoids sampled three months post-plating (**Fig. 1i**) confirmed that the organoids extend cellular processes embedding and surrounding the INFORM arrays. Immunostaining of CLARI-O-cleared organoids with neurofilament heavy chain (NF-H) confirmed the axonal identity of these processes (**Fig. 1j**), supporting structural integration forming through close contact with both top and bottom interfaces, which is essential for capturing electrophysiological activity with high signal-to-noise ratio (SNR).

Use of microelectrodes for stimulation confines current delivery to a spatially restricted region, minimizing spread to surrounding tissue and hence can enable selective recruitment of local neuronal populations^29^. Low electrode impedance is important to achieve high efficiency targeted microstimulation as well as increasing the SNR of single-unit and LFP recordings^30–32^. Thus, we deposited platinum nanoparticles (PtNP) onto the gold electrodes to lower their impedance from 1-2 MΩ to ∼50 kΩ (**Supplementary Fig. 7**). The spatial mapping of electrode impedances and electrochemical impedance spectroscopy (EIS) results show high uniformity across the interface (**Fig. 2a,b**) with an average impedance of 46.6 kΩ at 1 kHz. Cyclic voltammetry (CV) measurements confirmed characteristic platinum redox peaks and revealed that PtNP-coated electrodes exhibited greater charge storage capacity (CSC) and a wider water window than bare gold electrodes, enabling higher charge injection during microstimulation (**Fig. 2c, Supplementary Fig. 7**). Chronopotentiometry measurements verified that PtNP-coated electrodes sustained safe charge injection up to 125 μA (**Fig. 2d**) before the polarization voltage exceeded the water window (−0.9 to 0.9 V), confirming an expanded range compared to bare gold electrodes, which can only support 5 μA for safe stimulation (**Fig. 2d, Supplementary Fig. 8**). At the start of each weekly recording session, the interface channel impedances were measured. **Figure 2e** shows that the number of working channels (n = 6 interfaces, impedance < 2MΩ) remained above 90% and average impedances of the working channels remained stable over one year of in vitro recordings. Collectively, these results confirm that our technology exhibits the electrochemical properties required for safe targeted microstimulation and reliable longitudinal recordings.

**Figure 2.**
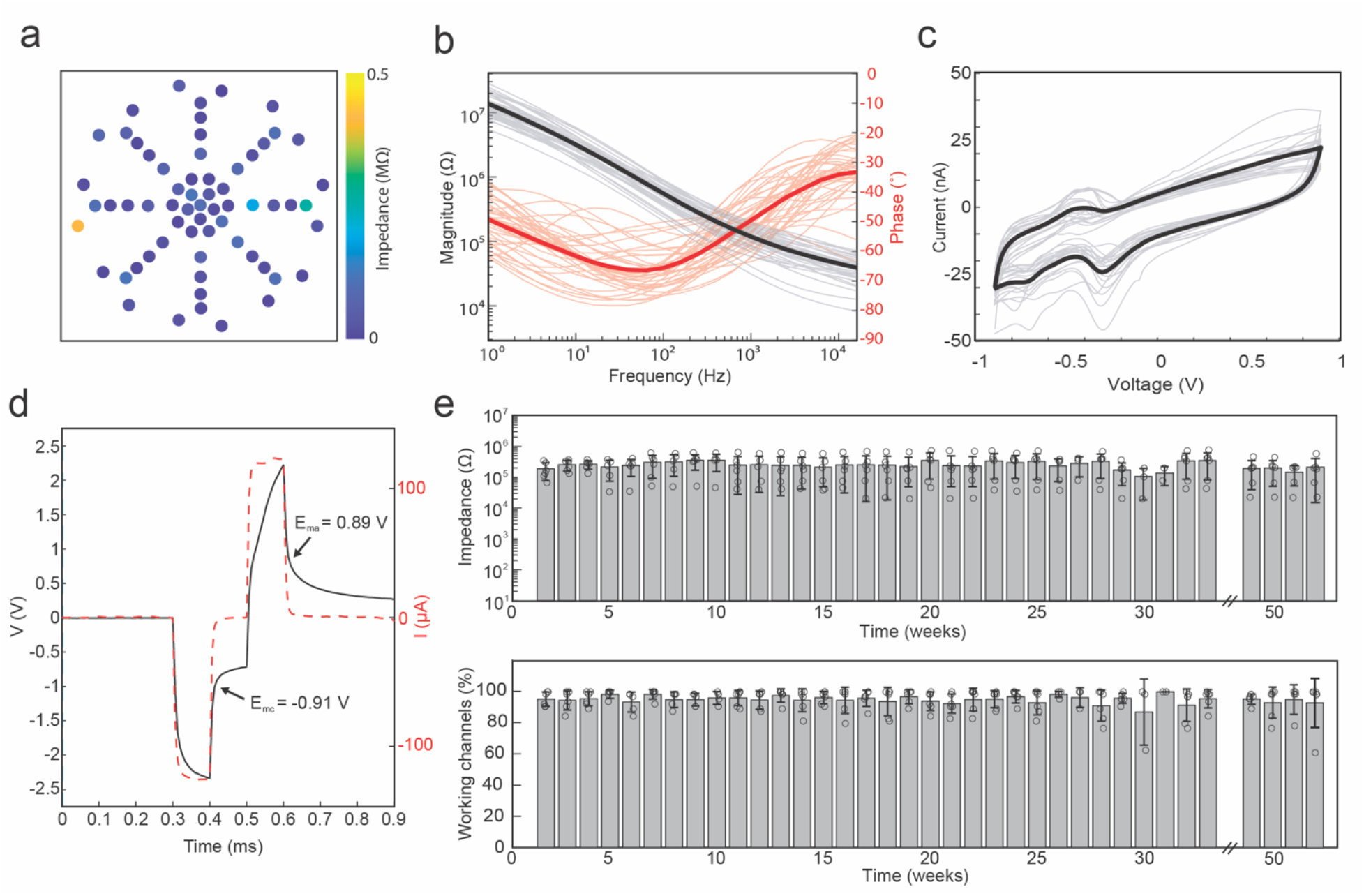
Electrochemical characterizations of the microelectrodes of the INFORM interfaces. (**a**) Impedances of 64 electrodes in a representative interface measured at 1 kHz with an average impedance of 46.6 ± 66.7 kΩ (mean ± SD). (**b**) EIS measurements after PtNP deposition; light traces are individual channels and dark marks the average of all electrodes (n = 40 electrodes from 2 interfaces). (**c**) Cyclic voltammetry results after PtNP deposition. Light traces are the averages of 3 cycles taken on individual channels and black trace marks the average of all electrodes (n = 19 electrodes from 2 interfaces). (**d**) Voltage transient generated using chronopotentiometry of a representative electrode after PtNP deposition. Current injected was 125 μA for 0.1 ms in cathodic then anodic pulses with a 0.1 ms period in between pulses. (**e**) Average impedance and percent of working channels (impedance < 2 MΩ) over 52 weeks in vitro (n = 6 interfaces). Circles indicate averages of individual interfaces and bars show mean ± standard deviation of all interfaces.

### Dorsal cortical organoids recapitulate cellular diversity and regional patterning

To characterize the cellular composition and regional identity of the organoids, we performed single-cell RNA sequencing (scRNA-seq) of 120 DIV cortical organoids derived from a male and female iPSC line. Based on the expression of canonical markers, we identified major cell types, including cycling progenitors, radial glia cells, intermediate progenitor cells (IPCs), astroglia, oligodendrocyte lineage cells (OLLCs), and distinct populations of excitatory neurons and inhibitory neurons (**Supplementary Fig 1a-b**). We observed a modestly expanded inhibitory neuronal population, that is consistent with our previous finding that exposure to neuronal activity medium (NAM), an activity-promoting medium, increases the representation of GABAergic interneurons in cortical organoids^33^. Despite differences in the abundance of cell types between two lines, all cell populations and subpopulations were recapitulated in organoids from both iPSC lines (**Supplementary Fig. 1c**). Our transcriptomic findings were further validated with cell-type specific nuclear markers for neurons (NeuN), astroglia (SOX9), and oligodendrocyte lineage (OLIG2) cells at an earlier 126 DIV timepoint as well as later 188 DIV timepoint (**Supplementary Fig. 1d**). Moreover, while organoids do not display the classical cortical lamination, they retain a small population of SOX2+ progenitors and express deep (CTIP2) and superficial (SATB2) cortical layer markers, confirming the presence of the major dorsally-derived neural cell types. Specific regional patterning of these organoids was transcriptomically verified with dorsal-lineage specific markers *PAX6* (progenitors) and *NEUROD6* (neurons) (**Supplementary Fig. 1f)**, Expression of *EMX1* (dorsal telencephalic) and *FOXG1* (forebrain) markers further confirmed dorsal forebrain identity, and the absence of ventral telencephalic (*NKX2-1, LHX6*), posterior/non-telencephalic (*HOXA2*), and non-neuronal mesenchymal (*COL1A1, PDGFRB*) and endothelial (*CD34*) markers (**Supplementary Fig. 1g)**. Overall, these results validated that the organoids largely differentiate into dorsal cortical neural cell types, consistent with the expected cellular features of organoid development.

### Simultaneous recordings of electrophysiology and cellular calcium activity reveal spatiotemporal network dynamics

Cortical organoids were transduced with GCaMP8s under a synapsin promoter (**Fig. 1h**, **Fig. 3a**) which allowed for simultaneous imaging of calcium transients alongside electrical recordings. Using two-photon microscopy in conjunction with electrophysiology, we investigated calcium activity patterns of large populations of neurons with cellular spatial resolution, while simultaneously recording electrical activity with high temporal resolution. Neurons within the organoids exhibited both sparse asynchronous firing as well as synchronous population bursts (**Supplementary Videos 2-4**). The low electrode impedance allowed us to record high-frequency activity including single neuron spikes and multi-unit activity (MUA) (0.5-4 kHz) as well as low-frequency local field potentials (LFP, 1-250 Hz). We observed a profound temporal alignment between calcium transients and increased spiking and amplitude of LFP peaks, indicating global events of coordinated neuronal activity within organoids (**Fig. 3b, Supplementary Video 1**). LFP was filtered into conventional electroencephalogram (EEG) frequency bands to further investigate the response. Significant correlation with the calcium peaks were observed with the signals in delta, theta, alpha, and spike bands (**Supplementary Figs. 9-10**). Highly synchronized bursts demonstrate that the organoid has enough excitable neurons and synaptic connectivity to generate recurrent population events^34^. Bursts can also suggest the presence of both inhibitory and excitatory neuron subtypes in the organoid networks (as shown in **Supplementary Fig. 1a-c)** and are characteristic of early cortical development in vivo^35–38^ Bursts are also indicative of synaptic strengthening in the brain through activity-dependent circuit-maturation during development^39, 40^. In fact, immunostaining at early 120 DIV and later 180 DIV timepoints show VGAT+ and VGLU1+ puncta identified adjacent to MAP2+ neuronal processes, signifying the presence of both inhibitory and excitatory synapses (**Supplementary Fig. 11)**. **Figure 3c** shows event-triggered spike waveforms detected across an organoid within a single recording session, where signals were localized to one or a few nearby electrodes which could be sorted into single- and multi-unit events (**Supplementary Fig. 12**). The firing pattern was observed to evolve over time, with electrodes showing both individual spontaneous spiking and coordinated population bursting activity at different time points. Burst rate progressed from undetectable at four weeks post-plating, when only individual spiking was observed, to 0.33-1 Hz one-year post-plating, reflecting the gradual change of synchronized network activity as organoid circuits developed and integrated with INFORM (**Supplementary Figs. 13-15**, n = 3 organoids). Because organoids were plated at 5 to 7 months of age, at which point synchronized network activity is expected to be present, we interpret this initial progression as reflecting the establishment of reliable electrode–neuronal coupling as the organoid anchored to and cellular processes extended along the interface, which led to the interface’s ability to capture changing organoid network dynamics over one year.

**Figure 3.**
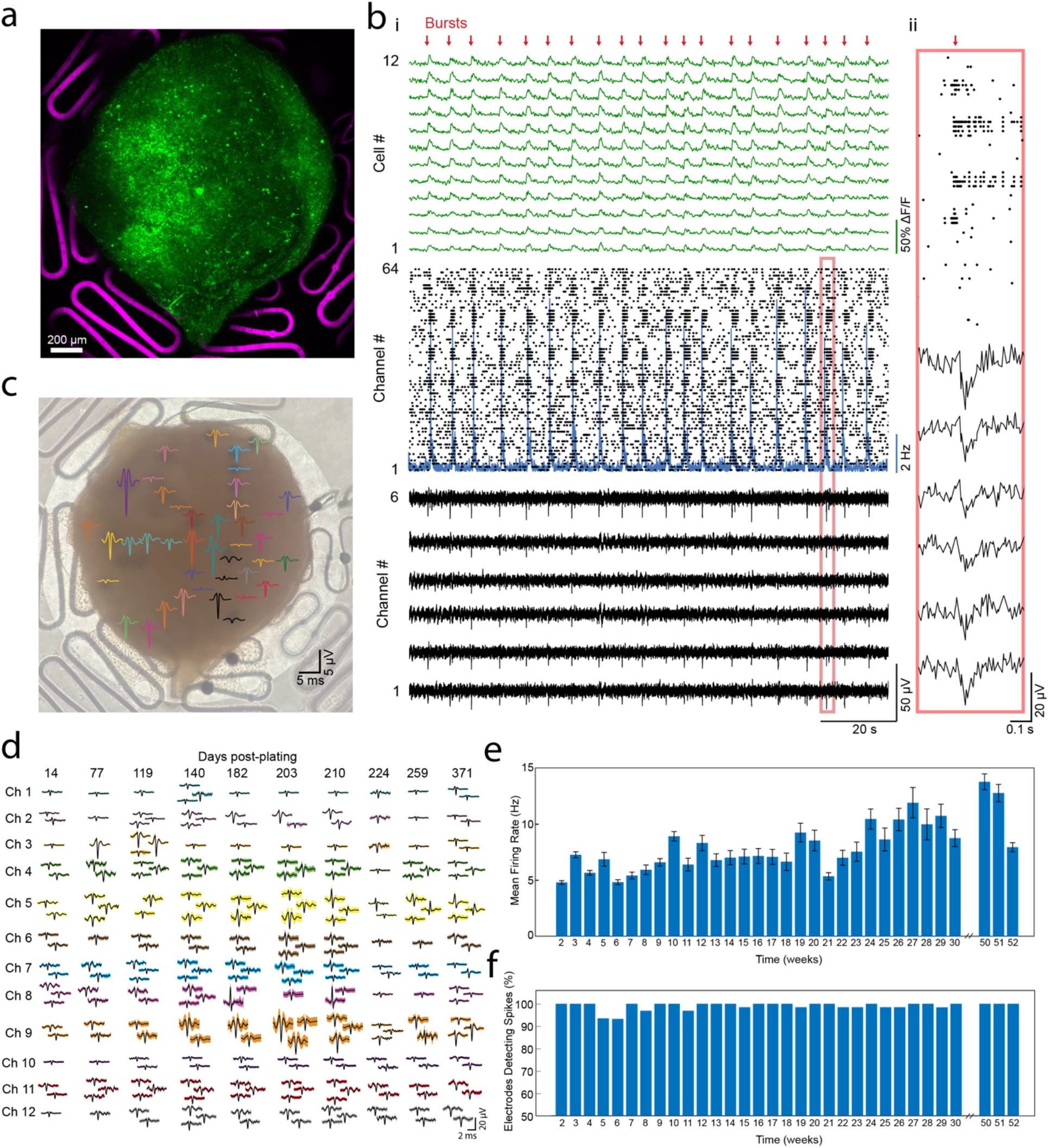
Multimodal longitudinal recordings of simultaneous electrophysiology and calcium dynamics. (**a**) Z-projection of a 260-day-old organoid (green) on an interface (magenta) 1.5-months post-plating. (**b**) (i) Multimodal experiments showing, calcium transients (top, n = 12 representative cells) simultaneously recorded with spikes (middle, n = 64 channels) and local field potentials of (bottom, n = 6 representative channels). Red arrows mark bursts where increases in calcium transients match increases in spiking activity and local field potential deflections. Organoid was 270 days old, 5 months post-plating. (ii) Callout showing a zoomed-in region of the LFP and spike raster marked in red box in (i). (**c**) Spatial representation of event-triggered spike waveforms detected in the organoid. (**d**) Sorted spikes detected on 12 channels over 371 days in vitro. (**e**) Mean firing rate of cells detected on channels for 52 weeks in vitro (n = 64 channels). Error bars indicate s.e.m. (**f**) Percentage of electrodes detecting spikes (on average, at least 3 spikes per second) (n = 64 channels).

To investigate the longevity of our interfaces, we recorded from organoids weekly for one year. Over this period, stable waveforms were recorded from the same electrode sites across sessions, with up to three distinct waveforms detected per electrode. Representative examples of detected spike waveforms distributed across different electrodes are shown in **Figure 3d**. The ability to consistently detect organoid spiking activity over an extended period of recordings provides a powerful means to monitor network stability and changes. Over time, spike waveforms slowly changed in shape and amplitude, likely due to neuronal maturation and migration, which caused different parts of neurons to be repositioned in closer contact with electrodes at different time points. The average spike rate generally increased over time, peaking around 50-60 weeks post-plating (n = 3 organoids, **Fig. 3e, Supplementary Fig. 16**), as organoid networks developed^1, 41^. Percentage of channels detecting spikes (≥3 spikes s⁻¹)^1, 26^ was maintained above 90% throughout multi-month recordings, indicating strong coupling between the interface and organoids enabling chronic recording stability (**Fig. 3f**). Overall, our platform surpasses the capabilities and performance of previous organoid interface technologies, achieving the highest percentage of channels detecting spikes, longest duration of activity tracking, and ability to continuously record local field potentials and spiking activity concurrently with cellular-resolution calcium imaging (**Supplementary Table 1**). These features position INFORM interface as a versatile technology to perform precise, long-term, spatiotemporal analysis of organoid networks.

### Assembloids show increased cellular synchrony over time

To investigate the dynamics of inter-organoid circuit formation, pairs of organoids were co-plated on individual interfaces and monitored longitudinally as they underwent fusion into assembloids (n = 6 organoids aged 5- to 7-months old on 3 interfaces, **Supplementary Fig. 17**). The organoids were observed to attach to one another through spontaneous fusion driven by cell migration, establishment of shared interface, and axonal outgrowth across the interface (**Fig. 4a,b**), with bridging likely mediated by Glial fibrillary acidic protein+ (GFAP+) astroglial cells and their processes, in line with the previously described processes accompanying organoids fusion^28, 42–44^. To assess the intra- and inter-organoid functional connectivity, we conducted weekly two-photon calcium imaging sessions (**Fig. 4b, Supplementary Video 6**). At two weeks post-plating, the organoids displayed internally synchronized bursting while remaining functionally independent from the other, as evidenced by uncorrelated activity across the pair (**Fig. 4c**, left). By week three, a subset of bursts became correlated across organoids, but most remained asynchronous (**Fig. 4c**, middle). By week four, activity across both organoids was almost entirely synchronized, indicating the establishment of widespread inter-organoid functional connectivity (**Fig. 4c**, right). Pairwise correlation analysis of calcium transients across all individual cells confirmed a progressive increase in inter-organoid correlation coefficients over time (**Fig. 4d**). Longitudinal quantification of the inter-organoid synchrony index^45^ over eleven weeks further demonstrated an increase in synchrony until the two organoids fired in near-perfect unison (**Fig. 4e**). Calculating the synchrony in two other assembloid pairs revealed that synchronization could even emerge before the two-week time point (**Supplementary Fig. 18**). High resolution two-photon imaging supported these findings through the observation of cells bridging the inter-organoid interface (**Fig. 4f**). To determine whether one organoid drove network-level activity in the fused pair, we examined the temporal onset of calcium peaks across cells with single-cell resolution. An average delay of 0.82 seconds was observed across the assembloid boundary (**Supplementary Fig. 19**), an example burst of which is shown in **Fig. 4g**, with cells in the left organoid (**Fig. 4g**, organoid 1) preceding those in the right (**Fig. 4g**, organoid 2). These results identified an apparent directional hierarchy in activity propagation, modeling long-range circuit formation observed in the developing brain^46–48^. These findings suggest directionality in activity propagation between the fused organoids and emphasize the dynamic, non-linear nature of assembloid circuit maturation, where critical connectivity transitions can emerge and resolve within days, underscoring the necessity and benefits of frequent longitudinal recording strategies.

**Figure 4.**
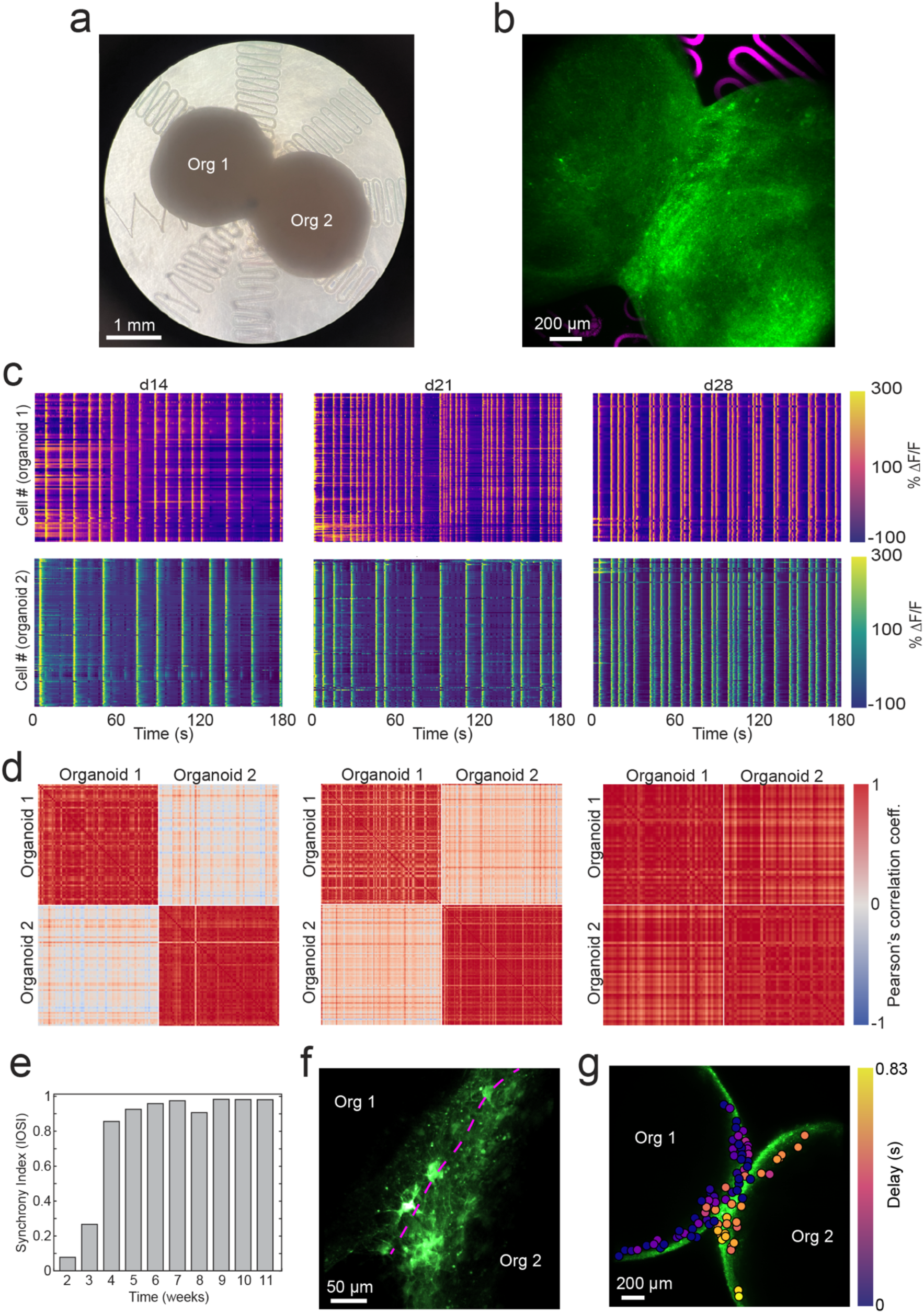
Assembloids show increased synchrony as functional connections form. (**a**) Brightfield image of two organoids plated in an assembloid configuration. Picture taken 11 weeks post-plating. (**b**) Two-photon z-projection of the same assembloid (green) on an interface (magenta) as in (a). (**c**) Heatmap of calcium transients of individual cells in organoid 1 and organoid 2 marked in (a) at two (left), three (middle), and four (right) weeks post-plating. (**d**) Pearson correlation coefficient heatmaps showing correlation of the calcium transients of cells in (b) at two (left), three (middle), and four (right) weeks post-plating. (**e**) Inter-organoid synchrony index of the assembloid over eleven weeks. (**f**) Two-photon microscopy image of the assembloid interface at three weeks post-plating. (**g**) Delay map of the calcium peaks for cells in both organoids for a representative burst event; delay is relative to the time of the earliest cell calcium peak.

### Targeted microstimulation elicits responses in localized subnetworks within organoids

To move beyond correlational observations and to enable precise interrogation of network dynamics, bipolar microstimulation was delivered via two of the 128 available electrodes, with one electrode serving as the cathode delivering a negative-first cathodic-anodic biphasic pulse and the other as an anode delivering the opposite polarity pulse (**Fig. 5a**). The bipolar microstimulation configuration confines current flow within a localized dipole, generating steep electric field gradients that minimize current spread and excite cellular activity at substantially lower charge densities than single electrodes with a distant ground configuration^49, 50^. Systematic parameter optimization across stimulus amplitudes (20–100 µA) and pulse counts (1–20) identified 100 µA and 10 pulses as the conditions eliciting the strongest calcium responses and were used for all subsequent experiments (n = 3 organoid interfaces, **Fig. 5b, Supplementary Fig. 20**). Microstimulation reliably evoked increased calcium burst frequency and elevated spiking activity (**Fig. 5c**), further quantified by averaging the spike band power (0.5-4 kHz) across microstimulation epochs (**Fig. 5di**). Critically, electrodes proximal to the microstimulation pair showed elevated responses, suggesting that focal microstimulation directly engaged cells close to the stimulating electrodes.

**Figure 5.**
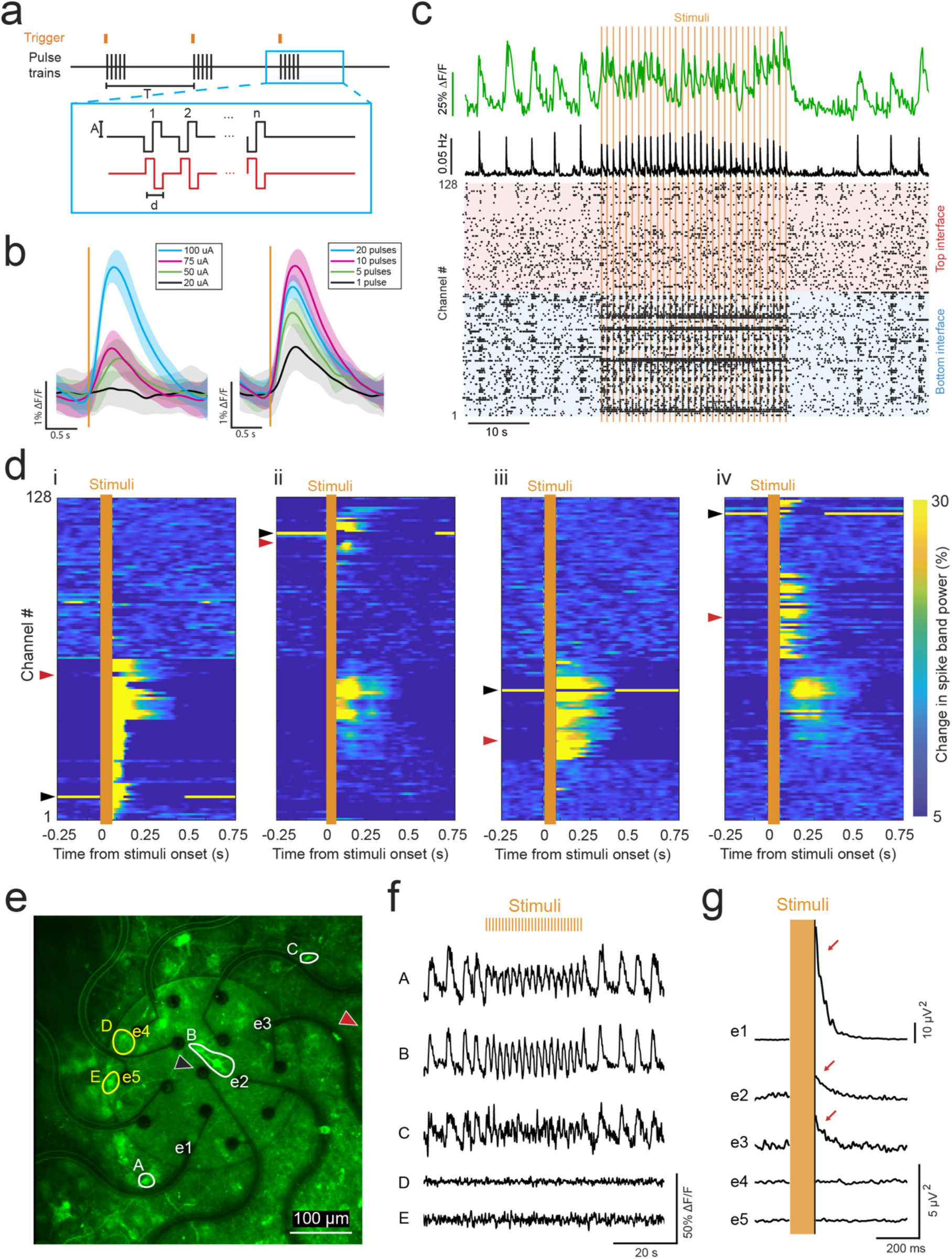
Electrical microstimulation of organoids. (**a**) Microstimulation configuration; two electrodes are chosen to deliver anodic-(red) or cathodic-first (black) biphasic pulses simultaneously. The pulses are defined by amplitude (A), pulse duration (d), and number of pulses (n) per trigger signal (orange). (**b**) Average calcium response taken of a whole organoid FOV in response to microstimulation of varying amplitude and number of pulse parameters (results shown as the mean ± standard deviation of 31 stimulations). (**c**) Calcium (green) and spiking activity response to microstimulation (orange); spike rate (middle) is the rolling time-averaged summation of individual spikes detected on all 128 channels (bottom) in the top (red) and bottom (blue) interfaces. (**d**) Average spike band power response in all 128 channels of the top and bottom interface to different microstimulation electrode pairings (n = 62 microstimulations); the black arrowhead marks the cathodic microstimulation channel and the red arrowhead marks the anodic microstimulation channel. Note panel i is generated using data from the same recording as c. (**e**) Two-photon microscopy image of cells underlying an interface. Letters mark cells and numbers mark electrode channels referred to in (f) and (g). Cells marked with white (A, B, C) responded to stimuli and cells marked with yellow (D, E) indicates non-active cells. Cathodic and anodic microstimulation channels are indicated by black and red arrowheads respectively (anodic microelectrode was slightly outside the frame). (**f**) Calcium response shows elevated activity for cells A, B, C while no activity for cells D and E. (**g**) Average spike power response to microstimulation shows increased power in electrodes (e1, e2, e3) close to active cells (A, B, C) while no power increase for electrodes (e4, e5) in vicinity of inactive cells (D, E) (n = 30 microstimulations).

To test whether distinct subnetworks could be recruited on demand, microstimulation electrode pairs were systematically chosen across varying locations on the interfaces. Notably, each new pairing (**Fig. 5di-iv**) selectively activated discrete subnetworks, some even seen in both top and bottom electrode interfaces, suggesting a spatially heterogeneous functional organization of subnetworks within the organoid. This spatial selectivity represents a marked advance over conventional approaches using non-spatially specific electrical stimulation^51, 52^ or full-field LED stimulation^10, 26^, which cannot isolate subregions of the organoid. Spatiotemporal comparison of calcium imaging and electrical recordings revealed a striking correspondence: cells exhibiting calcium transients (A, B, C in **Fig. 5e,f**) in response to microstimulation were spatially co-localized with electrodes (e1, e2, e3 in **Fig. 5e**) displaying elevated spike band power (**Fig. 5g**). In contrast, cells that do not show calcium transients (D, E in **Fig 5e,f**) also do not result in an increase in spike band power in the nearby electrodes (e4, e5 in **Fig. 5e,g**). This precise signal localization stems from the sub-cellular electrode geometry, which resolves high-frequency spikes from individual units rather than integrating activity across the neighboring neuronal cell population^53^. These results demonstrate high-resolution microstimulation and recording capabilities which enable systematic mapping of functional architecture in organoids.

### Electrical microstimulation reveals functional inhibitory subnetworks

Having observed subnetwork activation with targeted local microstimulation, we investigated functional responses at cellular resolution. Multimodal experiments involving microstimulation unmasked two functionally distinct subpopulations indistinguishable at baseline: an ’enhanced’ population reliably entraining calcium transients to the frequency of electrical stimulation, and a ’suppressed’ population exhibiting paradoxical silencing during electrical stimulation (**Fig. 6a-b, Supplementary Video 5**). These categories describe how a cell responded to stimulation and do not assign neuronal identity: a suppressed cell may be an excitatory neuron receiving an increased inhibitory input, whereas an enhanced cell may also be either an excitatory or inhibitory neuron, depending on the circuitry recruited by the stimulation. We hypothesized that the microstimulation activated presynaptic inhibitory and excitatory neurons projecting onto the observed cells, whose activity was captured within the two-photon imaging field of view. These enhanced and suppressed population dynamics mirror endogenous E/I mechanisms observed in vivo^54–56^, making their characterization in organoids important for validating their utility as disease models for disorders rooted in E/I imbalance such as epilepsy, Parkinson’s disease, and autism.

**Figure 6.**
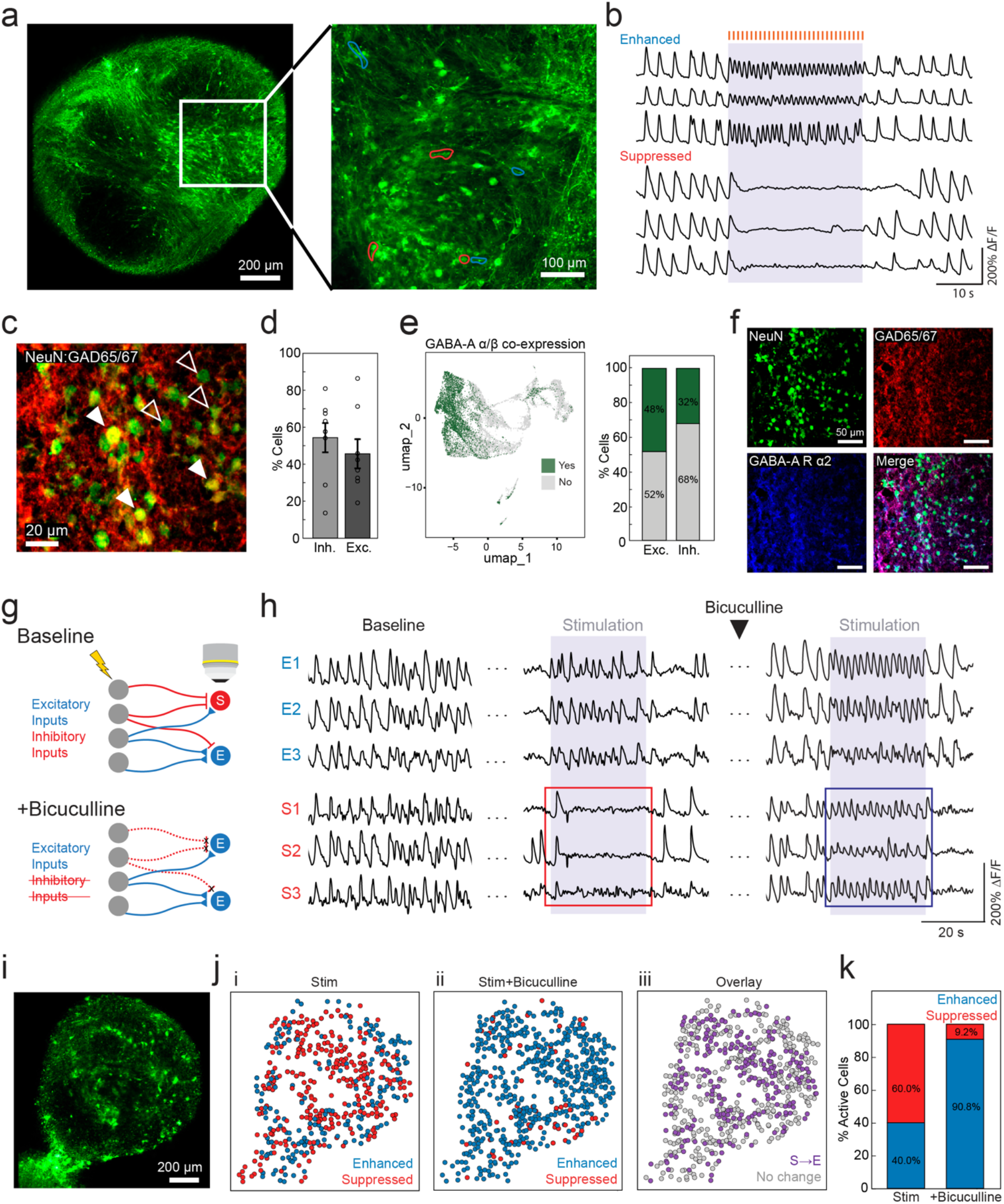
Spontaneous and stimulated calcium transients before and after bicuculline administration. (**a**) Two-photon microscopy image of a cellular network; blue outlines mark three representative cells that were enhanced by microstimulation and red outlines mark cells three representative cells that were suppressed by microstimulation. (**b**) Calcium transients of the cells marked in (a) during baseline and microstimulation (orange). (**c**) Overlap of NeuN and GAD65/67 indicate inhibitory cells (filled arrowheads) and NeuN+ cells not containing GAD65/67 (open arrowheads) are excitatory. (**d**) Quantification of inhibitory (Inh.) and excitatory (Exc.) cells in 180-day-old organoids shows 54.4 ± 20.9% inhibitory and 45.6 ± 20.9% excitatory neurons (n = 6 organoids derived from two cell lines). (**e**) Feature plot of GABA-A receptor α and β co-expression on the entire population. If it expresses at least one α subunit and at least one β subunit transcript, indicating capacity for the cell to function as a GABA-A receptor, the cell is colored green. Otherwise, the cell is colored gray. Bar graph quantifies 48% of excitatory and 32% of inhibitory neurons have at least one α and β subunit. (**f**) Immunostaining showing presence of GABAergic cell types and GABA-A α2 receptor. (**g**) Diagram of cells subject to microstimulation sending inputs to cells observed with calcium imaging. Addition of bicuculline disinhibits the network and prevents inhibitory inputs. Observed cells are labeled E (enhanced) and S (suppressed) according to their functional response. These cells may express excitatory or inhibitory outputs; the labels describe response category and make no claim about neuronal identity. (**h**) Calcium transients of three representative cells that were enhanced (E1-3) or suppressed (S1-3) by microstimulation during baseline (left), during microstimulation (middle), and during microstimulation after bicuculline was administered (right). Suppressed cells were inactive during initial microstimulation (red box) and then became responsive to microstimulation following bicuculline administration (blue box). Response was replicated in recording sessions of an organoid on days 562 and 632. (**i**) Two-photon microscopy image of the organoid plane used for analysis in (j-k). (**j**) Plots corresponding to the cells shown in (i) labeling cells as enhanced (blue) or suppressed (red) before (i) and after (ii) bicuculline was administered. (iii) marks cells that changed from suppressed to enhanced (purple) or experienced no change (gray) in their response to stimulation following bicuculline administration. (**k**) Quantification of the percent of enhanced and suppressed cells shown in (i-j) before and after bicuculline administration shows that 50.8% of cells changed from suppressed to enhanced (average percent change calculated from n = 120 microstimulations from two independent recordings from a representative organoid interface on different days).

To investigate the mechanisms behind the population-specific responses seen during electrical microstimulation, we validated the presence of inhibitory and excitatory neuron subpopulations (**Supplementary Fig. 1a-c, e)** and the intracellular machinery which may be driving this phenotype. Transcriptomically, subclustered inhibitory neuron populations demonstrated robust expression of canonical GABAergic neuron makers*, DLX5* (immature interneuron)*, GAD2* (pan-interneuron), and *SLC32A1* (functional interneuron) (**Supplementary Fig. 21a**). Immunostaining for the marker of mature neurons (NeuN+) colocalized with GAD65/67 identified GABAergic inhibitory interneurons (NeuN+/GAD65/67+), while NeuN+ cells with no detectable GAD65/67 expression were classified as excitatory neurons (**Fig. 6c**). Quantifications revealed that inhibitory neurons comprised 54.4% of the NeuN+ neuronal population (**Fig. 6d**), indicating a presence of inhibitory interneurons that can explain the strong suppressive response elicited by microstimulation (**Fig. 6b**). To validate the capacity to respond to inhibitory input, we first validated the expression of GABA-A receptor subunits at the mRNA level (**Supplementary Fig. 21b and Fig. 6e**). More specifically, both inhibitory and excitatory neurons express multiple GABA-A receptor subunits spanning developmental, core, and mature receptor composition (**Supplementary Fig 21b)**. Cells with functional GABA-A receptors were defined as ones co-expressing α and β subunits^57^ with co-expressing cell populations comprised of 48% of excitatory neurons, and 32% of inhibitory neurons **Fig. 6e**. Immunostaining confirmed the expression of GABA-A receptor α2 subunit at the protein level in both excitatory and inhibitory neurons, thus verifying the presence of cellular machinery for processing GABAergic input (**Fig. 6f**). Supported by immunostaining and transcriptomic data, the detection of suppressed cells during stimulation suggests that a sufficiently strong inhibitory synaptic input may override excitatory drive, via spatiotemporal summation of GABAergic inhibitory inputs from the stimulated population (**Fig. 6g**). In contrast, in enhanced cells, excitatory inputs outweigh inhibitory drive, leading to entrainment of the spiking activity to the stimulation frequency (**Fig. 6g**). The presence of inhibitory neurons and the fact that we used a high-frequency (100 Hz) microstimulation paradigm which preferentially stimulates inhibitory neurons^54, 58–61^ further supports that the activation of inhibitory synaptic drive resulted in the observed enhanced and suppressed population response patterns.

To further investigate the preferential functional responses of neurons to microstimulation, and to determine whether the stimulation was causing suppression through synaptic inhibition or another source such as depolarization block, we administered bicuculline (10 μM), a pan-GABA-A receptor competitive antagonist, which reduces chloride channel opening and diminishes inhibitory inputs (**Fig. 6g**).

Under baseline (no stimulation) conditions, the enhanced vs. suppressed populations were indistinguishable (**Fig. 6h**, left panel), while stimulation resulted in distinct responses between enhanced and suppressed populations (**Fig. 6h**, middle panel). In contrast, administration of bicuculline preserved activation of enhanced cells and, crucially, converted previously suppressed cells to an enhanced state, confirming the role of GABAergic inhibition in mediating stimulus-evoked suppression (**Fig. 6h**, blue and red boxes). These findings suggest that the stimulus-evoked suppression was mediated by GABA-A receptor activation, resembling the process by which GABAergic interneurons gate neuronal firing in vivo, a canonical inhibitory circuit motif of the central nervous system^54, 62^. Bicuculline-mediated GABA-A receptor blockade disinhibited the stimulation-dependent suppression (**Fig. 6h**, right panel), allowing excitatory inputs to elicit action potentials in the imaged cells^40, 63^.

We then characterized the functional inhibitory effects on the entire network by quantifying the change in cell responses during stimulation before and after bicuculline addition (**Fig. 6i-k, Supplementary Fig. 22**). While bicuculline addition only slightly increased overall baseline activity (**Supplementary Fig. 23**), adding stimulation revealed a far more dramatic change. Before bicuculline, 60% of active cells in the FOV were suppressed and 40% were enhanced during stimulation. Following bicuculline addition, the proportion of suppressed cells dropped to 9.2% while the proportion of enhanced cells rose to 90.8% (**Fig. 6k, n = 2 recording sessions from a representative organoid interface. Separate sessions’ results shown in Supplementary Fig. 24**). This represents a net shift of 50.8% of cells switching from suppression to enhancement upon bicuculline addition, suggesting that these populations were sensitive to GABA-A inhibition, consistent with microstimulation recruiting inhibitory circuitry. Once bicuculline blocked the GABAergic inputs to those cells, domination of excitatory inputs caused the cells to become enhanced during microstimulation. This interpretation is further supported by our immunostaining and transcriptomic results showing the presence of GABA-A receptor machinery in both excitatory and inhibitory neurons (**Fig. 6e-f**, **Supplementary Fig. 21**). The neurons which remained suppressed after bicuculline could be suppressed through mechanisms independent of GABA-A-receptor-mediated inhibition as bicuculline administration did not alter their response to stimuli. These results show that multimodal experiments enabled by our INFORM interface can probe functional neural populations at fine spatiotemporal scale to reveal inhibitory subnetworks not engaged in passive recordings.

Furthermore, we assessed the spatial distribution of enhanced and suppressed neurons and found that they were uniformly distributed across the organoid (**Fig. 6i,j**). This spatial profile suggests that their responses to microstimulation were indirect and mediated through network-level connectivity rather than direct activation through localized stimulation, which preferentially activates cells close to the site of microstimulation delivery (**Fig. 5d**). We investigated whether varying the location of microstimulation electrodes could controllably elicit enhancement and suppression on distinct subpopulations, a feature essential for biological computing or neuromodulation tasks, where precise, varied inputs must be delivered to activate specific cell populations. We found that indeed, different microstimulation configurations resulted in different subpopulation responses (**Supplementary Fig. 25**), demonstrating that by leveraging various electrode pair configurations, our INFORM interface can selectively and differentially engage functional subnetworks in cortical organoids.

## Discussion

We demonstrated INFORM, a flexible 3D interface platform that enables longitudinal, multimodal interrogation of cortical organoid networks at single-cell resolution. Conformal wrapping of 128 electrodes across both organoid hemispheres, consisting of 2-µm-wide metal traces and 12-µm electrode openings, achieved dense electrophysiological coverage for recording and stimulation while preserving full optical access for simultaneous two-photon calcium imaging. The interface geometry and PDMS housing were designed together so that four modalities: electrophysiology, targeted microstimulation, two-photon imaging, and pharmacological intervention could be applied to the same intact organoid simultaneously. While use of subsets of these modalities has precedent, the simultaneous application of all four is the first such demonstration in organoid systems. The PDMS well is paired with a glass window that prevents evaporation from heated media during recordings, allowing a 10x air objective to be used without lens condensation and avoiding the contamination risk of lowering an immersion objective into the culture week after week, while side ports permit delivery of pharmacological agents and lateral contact pads allow electrical connection that is removable between experiments and outside of the optical path. These design choices were essential to combining multiple modalities and led to comprehensive characterization of organoid networks by bridging the benefits of each.

Long-term reliability is a well-known limitation of flexible polymer arrays, and previous organoid interfaces have reported recordings spanning only a single day to a few months^7–11^. Serpentine arms of minimal thickness (4 µm) and width (10–34 µm) made the interfaces maximally compliant, allowing them to conform to organoids of any size or shape rather than requiring a predefined geometry^8, 9^, and yielded stable electrode impedances and above 90% channel yield throughout the year in culture, the longest continuous organoid recording reported to date. Targeted bipolar stimulation combined with multimodal readouts revealed functional subnetwork dynamics on the single-cellular scale not achievable with passive or unimodal recordings. Stimulation selectively engaged distinct neuronal subpopulations, and addition of bicuculline confirmed that stimulus-evoked suppression was mediated by GABAergic inputs gating spinking activity, with receptor blockade shifting neuronal responses from suppression towards enhancement, consistent with disinhibition. Critically, the scope of inhibitory input influence on neuronal populations was uncovered using stimulation, underscoring the necessity of active perturbation for comprehensive circuit characterization.

Weekly recording of the organoid fusion into assembloids demonstrated the value of the INFORM interface for studying inter-regional circuit formation, a process whose rapid developmental transitions would be missed by less frequent recording strategies. As the fetal brain matures, long-range cortico–cortical connectivity emerges alongside increasing interhemispheric synchrony^48^. In line with this, progressive synchronization observed within assembloids may reflect the development of cortico–cortical projection neurons and the establishment of long-range functional connections that has been shown to begin forming in the organoids after two months in culture^23^. In this study we plated organoids at later developmental timepoints (5-7 months old), focusing the analysis on functional connectivity between the already established networks rather than early network assemblies. Future studies plating younger organoids paired with our ability to record organoids for over one year provides access to developmental stages that enables monitoring of the progression toward mature, long-range functional circuitry at a comprehensive temporal resolution.

In addition, continuous single-cell resolution measurements enable tracking of individual neuronal activity patterns and stimulus responsiveness across developmental timepoints. This offers a clearer and more precise strategy than inferring dynamics from cross-sectional snapshots or post-fixation immunostaining. The combination of electrophysiological and optical functional readouts allows direct alignment of single-cell activity with network-level dynamics and is particularly valuable in organoid systems, where cellular maturation trajectories are still developing.

Future applications of this platform include investigation of other organoid and assembloid types, including studying circuit dynamics between different brain regions, as the INFORM interface is not constrained to a specific organoid size. Interrogation of E/I subnetworks in organoids could provide insight into mechanisms of cortical computation such as gamma oscillations, network synchrony, and propagating waves thought to encode spatiotemporal and predictive information^12–15^. INFORM interface’s high resolution recording and stimulation capabilities position the platform as a powerful neuromodulation tool to provide therapeutic stimulation to organoids modeling neurological diseases mediated by neuronal excitability such as epilepsy and Parkinson’s Disease^64–66^, train organoids for biological computing tasks^67^, or guide functional maturation^68–70^. The effects of microstimulation on excitatory and inhibitory cells remains elusive despite being fundamental to numerous in vivo neuromodulation therapies including deep brain stimulation^61, 71^, intracortical microstimulation^72, 73^, and brain-computer interfaces^58, 74^. Using this platform to study how microstimulation affects the balance of E/I subnetworks in organoids could accelerate the development and optimization of these therapies, benefitting millions of people suffering from neurological disorders globally. Finally, optical compatibility with two-photon imaging further extends the platform to optogenetics-based selective activation of GABAergic or glutamatergic cells during subnetwork studies^75^. Similarly, labeling excitatory and inhibitory cells with specific calcium indicators, such as ones under DLX or CaMKII promoters^76, 77^ was out of the scope of this study but in the future, would resolve in real-time exactly which type cells are being affected by the microstimulation to better elucidate the suppressed and enhanced subnetwork dynamics demonstrated here. Specifically, since the stimulation was delivered at 100 Hz frequency, which has been shown to preferentially activate inhibitory cells in literature, it would be interesting to see if the enhanced cells were primarily inhibitory or excitatory cell types and experiment with other frequencies to observe subsequent effect on network activity in the organoid system. The vast capabilities and excellent spatiotemporal resolution of our organoid interface establish it as a holistic framework for investigating human cortical network development, disease-related circuit dysfunction, and pharmacological responses in vitro.

## Methods

### COMSOL simulations

To evaluate the mechanical behavior and -z directional deflection of the flexible interface architectures, theoretical simulations were performed using the plate physics interface in COMSOL Multiphysics. Two-dimensional geometric models of the interfaces, featuring a central orbital electrode arrangement and varying numbers of serpentine arms (8, 16, and 32), were imported from AutoCAD DXF files and assigned a uniform thickness of 4 µm. Since a Parylene-C material profile is natively unavailable, the interfaces were modeled as linear elastic materials using a modified polyethylene terephthalate (PET) base template. Core mechanical properties for Parylene-C were manually defined utilizing established literature values, specifically a density of 1289 kg/m³, a Young’s modulus of 4.0 GPa, and a Poisson’s ratio of 0.4^78, 79^. During the simulation, rigid boundary conditions were applied to the external perimeter of the interface to simulate physical anchoring. To reflect the in vitro aqueous environment, a 95% reduction to the nominal 1 mg organoid mass was approximated to account for the buoyant force exerted by the surrounding culture media, exhibiting a density ∼1041 mg/cm³ comparable to water^7, 80, 81^. The organoid was represented as an applied centered boundary face load, corresponding to an effective mass of 0.05 mg and a boundary load of ∼2.5 Pa over the 500 µm diameter contact region. Simulations were performed at the design stage to test relative deflection of serpentine geometries under organoid weight, thus no constitutive material model was assigned to the organoid and its compliance and deformation were not simulated. Finally, the models were meshed and solved to extract theoretical minimum and maximum deflection values, generate 3D spatial deflection gradient heat maps, and plot Von Mises stress distributions to identify regions of high-stress concentration across the varying serpentine arm configurations.

### Fabrication of the INFORM interface

Fabrication of the INFORM interface consists of the following steps (**Supplementary Fig. 2b**). Four-inch silicon wafers were spun with PMGI SF3 as a sacrificial layer, baked at 125°C for 5 min, and then a two-micrometer layer of parylene C was deposited onto the wafer. Ten nanometers of chromium and 100 nm of gold were sputtered onto the wafer using Denton 18 Discovery System. The metal wires and contact pads were patterned using photolithography and wet etching (Gold Etchant TFA, Chromium Etchant 1020AC). A 10 μm encapsulation layer of Parylene C was then deposited. A photoresist photomask was patterned to uncover the 1-cm-diameter interface arm area and 8 μm of parylene C was etched using oxygen plasma so the interface arms were encapsulated between two 2-μm thick layers of parylene C but the rest of the interface remained 12-μm thick for easier handling and support. Then, the electrode openings and contact pads were etched using photolithography and oxygen plasma. Next, Al_2_O_3_ was sputtered onto the wafers and patterned into a mask for the profile etch using photolithography and etching in buffered oxide etch (BOE). The wafer was diced with one interface on each wafer piece and the parylene C interface arm profile was etched using oxygen plasma. After profile etching, the interfaces were placed into BOE for 20 seconds which removed the Al_2_O_3_ mask and released the interface from the wafer. Using a tweezer to hold the interface against the wafer piece to prevent the thin interface arms from tangling in air, the interfaces were transferred to two DI water baths to rinse BOE then the sacrificial PMGI SF3 layer was removed through successive baths in AZ NMP, acetone, and IPA. Finally, the interface was scooped up using a wafer and, while still wet with IPA, a piece of PVA tape was gently placed onto the interface arm section to keep the arms from tangling during successive handling. The interfaces were placed into a petri dish and allowed to dry overnight before assembly with PDMS wells.

### Assembly of interfaces within PDMS wells

Kapton tape was placed onto the back of the connector portion of the interfaces to reinforce the contact pads. The PDMS wells consist of several layers of PDMS structures sandwiching the interfaces, on top of a 25×75 mm glass slide. The PDMS layers were made by pouring uncured PDMS (Silgard 184) into 3D printed molds and baking at 80°C for 2.5 hours. The first PDMS layer was glued to a 25×75 mm glass slide using uncured PDMS and baked in an oven at 80°C for 10 minutes. Then the interface was placed on top of the first PDMS layer and the second layer of PDMS was placed on top of the interface. Note, the connector portion of the interface protruded out of the side of the PDMS well for zero-insertion-force (ZIF) connector access. The sides were then sealed with uncured PDMS and the assembled device was baked at 80°C for 2.5 hours to fully cure.

### Electrochemical characterization

To characterize the interfaces, 0.01 M phosphate buffered saline (PBS, Sigma-Aldrich P3813 dry powder dissolved in deionized water) was added to the PDMS well of assembled interface devices, which allowed the PVA tape to gently detach from the interface arms. Gamry 600 Plus potentiostat was used for electrochemical characterizations. Electrochemical Impedance Spectroscopy, Cyclic Voltammetry, and Chronopotentiometry measurements were performed in PBS using a three-electrode setup with an Ag/AgCl wire as reference and platinum wire as the counter electrode. CV voltage was swept with a rate of 150 mV/s. The system was enclosed in a self-made faraday cage to eliminate noise. NanoZ (White Matter LLC) was used for electrodeposition of platinum nanoparticles (PtNP). The interface was submerged in a H_2_PtCl_6_ (0.05 M) and K_2_HPO_4_ (0.01 M) solution with a platinum wire as the counter electrode and a current of -25 nA was delivered between the working and counter electrode to deposit PtNP until impedances reached ∼50 kΩ.

### Cortical organoid generation

Cortical organoids (CO) were generated and cultured using a modified published protocol^27^. CO generation was done using a female WC-24-02-DS-B and a male DS1-iPS4-disomic cell-lines with normal karyotype^33, 82^. Once about 65% confluent, a 35-mm well of iPSCs was washed with 1 mL of PBS and dissociated using 1 mL of Accutase (cat. 7920, STEMCELL Technologies). It was incubated for 5 minutes at 37°C, and with the addition of 5mL of DMEM/F12 (cat. 11330–057, ThermoFisher Scientific, Waltham, MA, USA), homogenized into a single-cell suspension. Once centrifuged and resuspended in 1mL mTeST plus with 50 µM of Rock Inhibitor (cat. 12-541-0, Fisher Scientific, Pittsburgh, PA, USA), cells were plated in a low adherence V-bottom 96 well plates (cat. MS-9096VZ, S-Bio Prime, Constantine, MI, USA), at a density of 15,000 cells/well/150 µL of media. On day one, a media half-change (100 µL) was made with TeSR-E6 (cat. 0596, STEMCELL Technologies) supplemented with 2.5 µM of Dorsmorphin (cat. P5499, Millipore Sigma, Burlington, MA, USA) and 10 µM SB-431542 (cat. S4317, Millipore Sigma). This was repeated daily on days 3-6. Beginning on day 7, media composition was switched to a basal medium [Neurobasal A Medium (cat.10888022, Life Technologies, Carlsbad, CA, USA) containing B-27 supplement without vitamin A (1:50; cat. 12587, Life Technologies), GlutaMax (1:100, cat. 35050061, Life Technologies), 100 U/mL Penicillin/Streptomycin (1:100, cat. 15140122, Life Technologies) and Primocin (1:500; cat. ant-pm-1, Invivogen, San Diego, CA, USA)] and supplemented with fibroblast growth factor 2 (FGF2; 20 ng/mL, cat. 233-FB-25/CF, R&D Systems, Minneapolis, MN, USA) and epidermal growth factor (EGF; 20 ng/mL, cat. 236-EG-200, R&D Systems). COs were transferred to 24-well ultra-low attachment plates (cat. 07-200-602, ThermoFisher Scientific) on day 18 and cultured with one CO per well, at this point media changes were conducted every other day. Between days 25 and 39, media composition changes to basal media supplemented with brain-derived neurotrophic factor (BDNF; 20 ng/mL, cat. AF-450-02, PeproTech, Waltham, MA, USA), neurotrophic factor 3 (NT3; 20 ng/mL, cat. 450-03, PeproTech) and 1% Geltrex (cat. A1569601, Life Technologies). From day 40 onwards, COs were maintained in basal media with 1% Geltrex until transition to a neuronal activity medium NAM, as described previously^33^ [BrainPhys Neuronal Medium (cat. 05790, Stemcell Technologies) supplemented with B-27 supplement (1:50, cat. 17504044, ThermoFisher Scientific), N2 (1:100, cat. 17502048, ThermoFisher Scientific), 10 mM 20% d-glucose (cat. G7021, Millipore Sigma), lipid concentrate (1:500, cat. 11905031, ThermoFisher Scientific), 56.779 mM ascorbic acid (cat. 4403, Millipore Sigma), Penicillin/Streptomycin (1:100, cat. 15070063, ThermoFisher Scientific), and Matrigel (1:100, cat. 356234, Corning)] starting at day 90. Media composition was adjusted in increments of 20% for 5 days, until composition reached 100% neuronal maintenance media. COs were then maintained in neuronal maintenance media until functional experiments took place and throughout the duration of experiments.

### Viral infection

For live-cell calcium imaging, COs were transduced at day 120 using a GCaMP8s lentivirus vector under a synapsin promoter to target neuronal cells. Product information is summarized in Table 1. For transduction, COs were moved to a 1.5 mL tube and incubated for 30 minutes at 37°C in 30 μL of maintenance media with 2 μL of LV. After 30 minutes, 300 μL of maintenance media was added to the tube and COs were incubated overnight at 37°C. After at least 16 hours COs were moved back to maintenance wells with 800 μL of media. Fluorescent imaging could take place starting at 14 days post-infection.

**Table 1.**

| Virus |  | Catalogue Number | Distributor |
| --- | --- | --- | --- |
| Neuronal calcium indicator | pLV<sup>94</sup>-<br>PuroSYN1.jGCaMP8s | VB240904-1449tnm | Vector Builder |
| General reporter | pLV<sup>94</sup>-<br>BsdEF1A>mScarlet3 | VB240919-1466xpw | Vector Builder |

### Organoid plating on INFORM interfaces

After interfaces were assembled into the PDMS wells, the devices were cleaned through 1-hour baths in 70% ethanol then sterile DI water in the biosafety cabinet with the UV light on. After removing from DI water, the external portion of the device was dried and a 1% laminin solution (Sigma-Aldrich L2020) in BrainPhys Neuronal Medium (Stemcell Technologies 05790) was added to the well through 10 half media changes and allowed to incubate overnight to coat the interface. The next day, the laminin solution was replaced with BrainPhys through six half media changes and a temporary PDMS ring (3 mm inner diameter, 5 mm outer diameter) was placed on the interface to help center the organoids during the initial weeks of adhesion. The organoids were placed onto the interfaces using 10 mL serological pipettes and cultured in NAM. After 2-3 weeks, once the organoids were observed to have anchored to the interfaces through cellular projections, the PDMS rings were removed. To place the second interface on top of organoids, freshly fabricated interfaces were cleaned in 1-hour ethanol and sterile DI water baths, transferred using a glass coverslip. Then the interface was scooped up with the coverslip and placed onto the organoid for contact with the top hemisphere. The glass slide was carefully removed and another layer of the PDMS well was glued with uncured PDMS to seal the top interface onto the device. When placing the top interfaces, another PDMS ring was used to hold the top interfaces in place against the organoid since moving the devices in and out of the incubator caused movement of the top interface which we wanted to minimize to promote adhesion of top interfaces to organoids. A 35 mm Mattek imaging dish lid was placed on top of the PDMS well to minimize evaporation and the whole device was housed in a ten-cm dish for sterile transport between incubator, biosafety cabinet, and recording setup.

### Simultaneous electrophysiology and two-photon microscopy

Recordings were conducted weekly starting at 2 weeks post-plating. Media in the organoid well was exchanged with BrainPhys Imaging Optimized Neuronal Media (STEMCELL Technologies 05796) through four half media changes for recordings and culture media was replenished the same way after recordings concluded each session. Each recording lasted less than one hour to minimize time the organoids spent outside the incubator. The interface devices were placed on a custom heated sample stage calibrated to maintain the organoids at 35°C and a glass coverslip (Fisher Scientific 12541019CA) was placed on top of the well to prevent media evaporation. Calcium activity recordings of GCaMP8s-labeled organoids were collected using an upright two-photon microscope (ThorLabs Bergamo II) equipped with a SpectraPhysics Insight X3 femtosecond laser tuned to 925 nm. A 10x apochromatic objective (ThorLabs TL10X-2P) was used to acquire high-resolution videos of calcium activity. Frames were acquired in resonant scanning mode at a frame rate of 30 Hz with a 512 × 512 pixel field of view, corresponding to a line rate of 8000 lines s⁻¹. The slow-scan axis was oriented in the y-direction relative to the imaging frame. Electrophysiology was simultaneously acquired using RHD2000 (Intan Technologies) for non-stimulation and RHS Stim/Recording System (Intan Technologies) for stimulation experiments. External platinum wires were placed in the side ports of the interface device as ground and reference during recordings. Custom PCBs routing a 64-channel ZIF to Omnetics connectors were used to connect the interfaces to Intan headstages. The ZIF connector allowed for temporary connections to the interface, so no PCBs or hardware were kept inside the incubator between recordings. The sampling rate was set to 20 or 30 kHz and data was imported to MATLAB (MathWorks) for analysis. Electrophysiology and imaging data were aligned using a square wave pulse generated by the imaging system corresponding to each imaging frame sent to an analog input in the Intan data acquisition board.

### Electrical microstimulation

The RHS Stim/Recording System (Intan Technologies) was used for microstimulation experiments. Two interface microelectrodes were chosen in the software to deliverer bipolar microstimulation, one delivering a cathodic-first biphasic current pulse and the other delivering a simultaneous anodic-first pulse. Biphasic current waveforms were chosen because the balance of anodic and cathodic phases decreases charge accumulation, thereby reducing tissue damage and electrode degradation^83, 84^. Parameters including current amplitude, pulse duration, and pulse repetition number and rate were set in the Intan RHX Data Acquisition Software. Each phase of the biphasic pulse was set to 100 μs duration, with no inter-phase interval, delivered at 100 Hz frequency. The amplitude of the pulses ranged from 20 to 100 μA and the pulses were repeated 1 to 20 times per trigger signal. The microstimulation trigger signals were controlled through user keyboard clicks spaced 1 second apart. Simultaneous electrophysiology was acquired using the same Intan system and calcium signal was acquired using ThorImage software (ThorLabs v4.3). Baseline recordings were taken directly before bicuculline administration. Then, 10 μM bicuculline (Alomone Labs B-136) was pipetted into the imaging media and recordings were conducted ∼10 minutes following drug administration. After experiments were conducted, organoids were rinsed through six half-media changes using fresh culture media to remove the bicuculline.

### Electrophysiology and calcium data analysis

Calcium traces of cell and organoid ROIs were extracted from tiff images using ImageJ2 (NIH, v2.9.0/1.54p) and exported as CSV files. All further calcium and electrophysiology analysis was performed in MATLAB (Mathworks, v2024a). Calcium traces were normalized by baseline fluorescence (mean of the signal) and smoothed using a 0.167-second gaussian filter. Electrophysiology was preprocessed to remove imaging artifacts using notch filters (6^th^-order butterworth) at the imaging frame rate and harmonics. Microstimulation times were set to NaN in recordings to avoid misclassification as organoid response and not affect standard deviation calculations for spike detection. Local field potentials were detected by filtering electrophysiology data through a zero-phase, band-pass 6^th^-order butterworth filter from 1-250 Hz (designfilt.m, filtfilt.m). Spiking activity including single units and multi-unit activity was extracted by filtering electrophysiology data through a 0.5-4 kHz zero-phase, band-pass 6^th^-order butterworth filter (designfilt.m, filtfilt.m). Spike power was calculated by squaring the signal then smoothing using a 80-ms gaussian filter (smoothdata.m). Spike times were detected by thresholding ±3.5 times the standard deviation of the spike signal using findpeaks.m. Spikes were sorted into distinct spike waveforms using PCA and k-means gaussian clustering. The inter-spike interval was calculated for each cluster and spike clusters that did not violate the refractory period of spike timing were classified as single-units and others were classified as multi-units. To determine population spike rate, the spike raster of all channels on an interface was summed across time and smoothed using a 10-ms gaussian filter (smoothdata.m). Statistical significance for correlation between the calcium peaks and electrophysiology bands was computed by shuffling the electrophysiology signal 10,000 times, computing the mean power of each frequency band or spike rate from 0.7 s before to 0.1 s after each calcium peak during a 3-minute recording, can calculating p-value as the number of times out of 10,000 shuffled cases where the mean power or spike rate exceeded the unshuffled case.

### Assembloid synchrony analysis

Calcium data was processed with Suite2P^85^ for automatic ROI detection and extraction of fluorescence traces (F), neuropil signals (Fneu), and ROI statistics. Only ROIs classified as cells by Suite2P were used for analysis. To classify ROIs into two separate organoids, a two-component Gaussian Mixture Model (GMM) was fit to ROI centroids (scikit-learn)^86^. When GMM labels disagreed with Mahalanobis distance, the closer cluster was used. The upper cluster was labeled Organoid A and the lower Organoid B. After identifying the clusters, the neuropil (0.7 * Fneu) was subtracted from each trace and the 8th percentile of each trace was used as the baseline. We computed ΔF/F and z-scored across time. Next, using these traces, the Pearson correlations between all ROI pairs was computed. Matrices were sorted by organoid, with Organoid A first then Organoid B. To measure calcium activity synchronization, Synchronization Cluster Analysis from FluoroSNNAP^45^ was used. Eigenvalues of the correlation matrix were computed and compared to 200 surrogates made by Amplitude Adjusted Fourier Transform (AAFT). AAFT keeps each trace’s amplitude and power spectrum but breaks timing between ROIs. For eigenvalues above surrogate mean plus 2 standard deviations, the Synchrony Index was calculated as SI = (lambda_observed − lambda_surrogate) / (M − lambda_surrogate), where M is the ROI count. The first eigenvalue’s SI is the network synchrony index. To measure synchronization limited to organoid A vs organoid B, correlations were accounted for only between ROIs from different organoids. The largest singular value was then calculated, normalized by sqrt(M_A × M_B) and tested against 200 AAFT surrogates applied to Organoid B only, keeping Organoid A unchanged. The Inter-Organoid Synchronization Index (IOSI) was calculated as (observed − surrogate_mean) / (1 − surrogate_mean).

### Assembloid cellular delay times analysis

The calcium onset of a cell was defined as the first frame at which its z-scored ΔF/F crossed a fixed threshold. Within each burst window, every ROI crossing the threshold at any point during the window contributes one onset, and its delay is that onset minus the earliest onset in the window. The activation span of a burst is the largest of these delays, and spans are reported as median and interquartile range across all bursts of a session.

### Enhanced/suppressed cell classification

Cells were classified as “enhanced” or “suppressed” based on their functional response during stimulation versus non-stimulation periods. For each cell, the area under curve (AUC) was computed of processed calcium traces using only positive values, normalized by the number of frames in each period. The mean of ΔF/F was computed during baseline and stimulation. The AUC ratio was AUC_stim divided by AUC_nostim. The mean ratio was mean_stim divided by mean_nostim. A combined ratio was computed as 0.7 × AUC_ratio + 0.3 × mean_ratio. Then, cells were classified using thresholds on the combined ratio. Next, to track cells’ activity shift between baseline and drug recordings within the same day, cells were matched by spatial proximity. Pairwise Euclidean distances were computed between cell centroids. Cells were matched using greedy nearest-neighbor assignment, accepting only matches within 20 pixels. After gaining insight on overlapping cell locations, scatter plots were generated comparing cell populations between two recording sessions. Cell locations were plotted with colors indicating response type (blue for enhanced, red for suppressed). Bar charts were created showing the proportion of enhanced, suppressed, and unaffected cells in each recording session. This allowed comparison of how drug treatment affected the distribution of response types across conditions. Lastly, to track how individual cells changed from baseline to drug conditions, matching cells were color-coded by their transition type and plotted. Cells that changed from suppressed to enhanced were color-coded purple and cells with no change were color-coded gray.

### Immunohistochemistry

For immunostaining, COs were washed with PBS and left in 4% paraformaldehyde (PFA) overnight for fixation. To cryoprotect, they were then moved to 30% sucrose until dropped. Once dropped, the COs were embedded in 30% sucrose/Optimal Cutting Temperature compound (OCT; Sakura Finetek, Torrance, CA, United States) at 60:40 ratio and sectioned at 12 μm. Slides were kept at -80°C until use. Sections were washed three times with 0.25% Triton X-100 (cat. CAS 9036-19-5, Sigma Aldrich) in PBS (PBS-T), blocked for 1 hour at room temperature (RT) in 5% normal donkey serum (cat. D966, Millipore Sigma) in 0.25% PBS-T. Next, the sections were incubated with the primary antibodies diluted in the blocking buffer at 4°C overnight. The next day, slides were washed three times with PBS-T (10 minutes), followed by an incubation with secondary antibodies (in blocking solution at 1:500) for an hour at RT. Slides were washed two times with PBS-T and once with PBS. They were then coverslipped with ProLong™ Gold Antifade Mountant with DAPI or without DAPI depending on experiment needs (cat #P36930 and P36931, Thermo Fisher Scientific, Waltham, MA, United States). The primary antibodies used are summarized in Table 2. Stained sections were imaged using spinning disk confocal microscope (BC43, Andor).

**Table 2.**
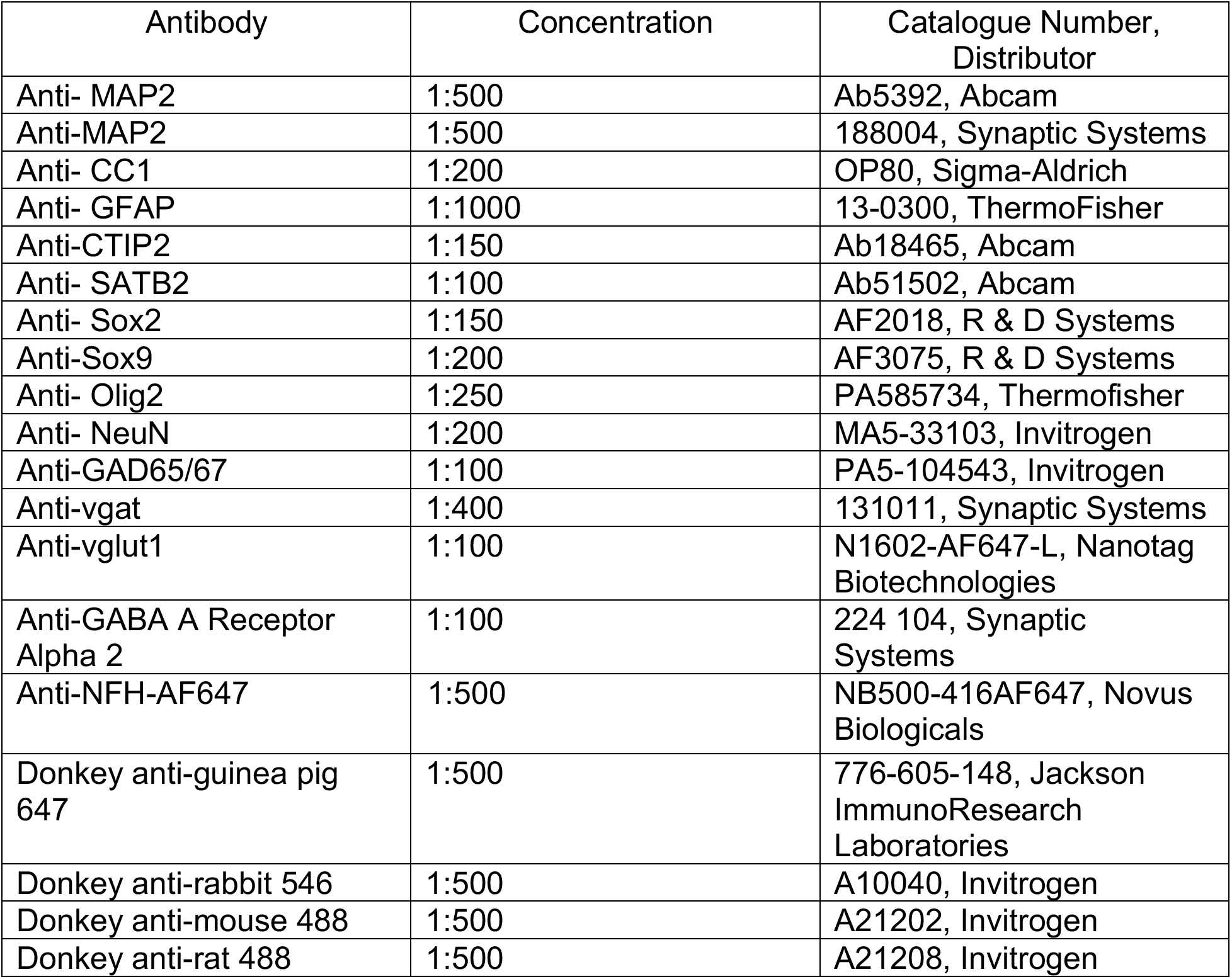
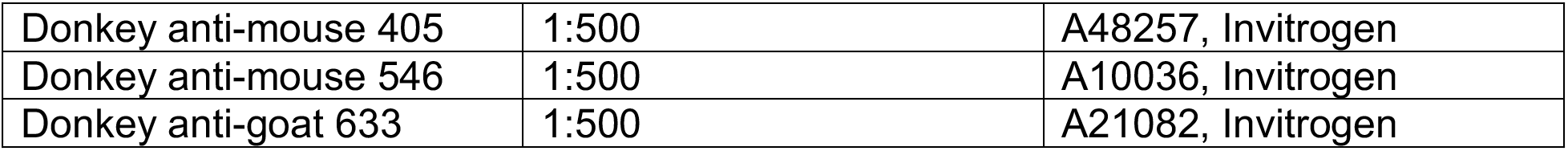

| Antibody | Concentration | Catalogue Number, Distributor |
| --- | --- | --- |
| Anti- MAP2 | 1:500 | Ab5392, Abcam |
| Anti-MAP2 | 1:500 | 188004, Synaptic Systems |
| Anti- CC1 | 1:200 | OP80, Sigma-Aldrich |
| Anti- GFAP | 1:1000 | 13-0300, ThermoFisher |
| Anti-CTIP2 | 1:150 | Ab18465, Abcam |
| Anti- SATB2 | 1:100 | Ab51502, Abcam |
| Anti- Sox2 | 1:150 | AF2018, R & D Systems |
| Anti-Sox9 | 1:200 | AF3075, R & D Systems |
| Anti- Olig2 | 1:250 | PA585734, Thermofisher |
| Anti- NeuN | 1:200 | MA5-33103, Invitrogen |
| Anti-GAD65/67 | 1:100 | PA5-104543, Invitrogen |
| Anti-vgat | 1:400 | 131011, Synaptic Systems |
| Anti-vglut1 | 1:100 | N1602-AF647-L, Nanotag Biotechnologies |
| Anti-GABA A Receptor Alpha 2 | 1:100 | 224 104, Synaptic Systems |
| Anti-NFH-AF647 | 1:500 | NB500-416AF647, Novus Biologicals |
| Donkey anti-guinea pig 647 | 1:500 | 776-605-148, Jackson ImmunoResearch Laboratories |
| Donkey anti-rabbit 546 | 1:500 | A10040, Invitrogen |
| Donkey anti-mouse 488 | 1:500 | A21202, Invitrogen |
| Donkey anti-rat 488 | 1:500 | A21208, Invitrogen |
| Donkey anti-mouse 405 | 1:500 | A48257, Invitrogen |
| Donkey anti-mouse 546 | 1:500 | A10036, Invitrogen |
| Donkey anti-goat 633 | 1:500 | A21082, Invitrogen |

### Organoid clearing with CLARI-O

CLARI-O was performed as previously described^28^. Briefly, organoids were fixed in 4% Paraformaldehyde for 48 hours. They were then washed at room temperature in 0.05 M TBS, pH 7.4, for 2hrs before being transferred into the CLARI-O solution (200 mM SDS containing 20 mM lithium hydroxide monohydrate, pH 9.0). Organoids were incubated at room temperature for 2 days, then transferred into a new solution and incubated at 37°C for 2 days. The CLARI-O solution was changed and organoids were incubated for another 2 days at room temperature on a rocker. The cleared tissue was then washed for 1 day at room temperature in 0.05 M TBS, pH 7.4 to remove the CLARI-O solution from the tissue, prior to staining and/or imaging.

### Imaging and image analysis through QuPath quantification

QuPath (V0.5.1) was used for neuronal sub-type quantification in cortical organoids. Processed and stained organoid sections were imaged using BC43 spinning disk confocal microscope (Oxford Instruments, High Wycombe, United Kingdom) and A 40X/NA1.3 oil objective. For the analysis, 2 organoid sections were used per organoid and for each section 2 randomly chosen fields of view (FOV) were imaged, with total of 4 FOV imaged and quantified per organoid. Organoids from two different lines were used and analyzed with QuPath using the “cell detection” function through the application of manual threshold and maintaining this threshold consistent across the images. GAD65/67+ cells were counted when colocalized with the neuronal maker, NeuN. Positive cells were labeled inhibitory, and negative cells were labeled excitatory. Measurements from 4 FOV were averaged per organoid and each organoid was considered as one biological replicate.

### Single cell RNA library preparation and sequencing

#### Single-cell dissociation

CO dissociation was performed at 121 DIV for differentiation 1 and at 123 DIV for differentiation 2. Two COs (1 from WC and DS1 lines) generated from different wells were pooled per sample and dissociated using the Worthingson Papain dissociation system (cat. LK003150, Worthington Biochemical Corp., Lakewood, NJ, United States) as previously described^87^.

Cell viability was assessed using Trypan-Blue and manual counting. Cell samples at a viability of at least 86% were used for processing with a target capture of 4000 cells/sample. Single-cell RNA sequencing was done according to 10X Genomics Chomium single cell system protocol. Sequencing was done in collaboration with the Single Cell Sequencing Core at Boston University School of Medicine.

#### Data Pre-processing and Quality Control

Sequencing data processing and quality control were performed as previously described using the 10X Genomics Cell Ranger V3 pipeline and Seurat package (v 5.0.0 and R version 4.4.0)^88, 89^. Cells with < 350 UMIs, Genes < 300, >20% mitochondrial reads were excluded. In addition, genes with low library complexity, defined by log10(genes to UMI ratio) < 0.8 were excluded. We normalized and scaled the data using NormalizeData and ScaleData functions. Dimensionality reduction (PCA) was then performed. To account for technical variation and inter-sample batch effects, Harmony^90^ was applied to the top 30 PCs using individual samples as the batch covariate. Harmony converged after 3 iterations and corrected embeddings were used for downstream analysis.

#### Cell Cluster Identification

Uniform Manifold Approximation and Projection (UMAP) dimensionality reduction was performed using harmony embeddings on normalized corrected assay. To identify clusters in reduced dimensional space, Louvain clustering (resolution 0.7, Seurat) was applied to Harmony reduced embeddings, resulting in 18 clusters. Cluster characterization was then performed using log-normalized and scaled RNA counts. Cell types were further annotated with canonical markers, resulting in 13 cell type groupings. Marker-based dotplots and heatmaps were generated in Seurat using scaled expression.

#### Inhibitory neuron lineage subclustering

Progenitor and inhibitory neuron clusters were subset from the full object and re-processed independently using the same workflow: normalization and scaling (NormalizeData, ScaleData), PCA, Harmony batch correction across samples (top 30 PCs), and UMAP on the Harmony-corrected embeddings. Clustering was re-run at resolution 0.4. Marker expression was visualized using feature plots on this re-embedded subset. Based on previous literature, *GAD2* and *SLC32A1* are canonical markers of GABAergic function, *PAX6* marks dorsal forebrain progenitors, *ERBB4* is associated with interneuron identity and *SCGN* marks a dorsally/CGE-associated cortical interneuron population^88, 91, 92^.

#### GABA-A receptor expression analysis

GABA-A receptor subunit genes were grouped by their reported developmental expression associations into developmental (*GABRA2, GABRA3, GABRA5, GABRB3*), core/structurally necessary (*GABRB2, GABRG2*), mature (*GABRA1, GABRD*) and other (*GABRB1, GABRA4, GABRA6, GABRG1, GABRG3, GABRE, GABRQ, GABRR2*) categories^89, 93^. Excitatory ((Newborn EXN, EXN deep layer 5-6, EXN deep layer 4-6, EXN superficial 2-4, Unspecified EXN) and inhibitory neurons (InN_1 and immature InN) were then subclustered via previous annotation. Log-normalized subunit expression was visualized across the subclustered groups as violin plots. To assess the capacity to assemble a functional GABA-A receptor, each cell was scored as positive if it expressed at least one α-subunit and at least one β-subunit transcript in the raw counts, and negative otherwise; this binary score was projected onto the UMAP. Under this criterion, 32% of inhibitory cells and 48% of excitatory neurons were positive.

## Acknowledgements

This research was supported by grants from NIH (DP2 EB030992) to D.K., (U01 NS139877) to D.K. and E.C., grants from NIH/NIA (R21AG080269) to E.Z. and (RF1AG088529) to M.T and E.Z. Fabrication of the INFORM interface was performed at the San Diego Nanotechnology Infrastructure (SDNI) of University of California San Diego, a member of the National Nanotechnology Coordinated Infrastructure, which is supported by the National Science Foundation (grant ECCS-1542148).

## Author Contributions

This work was conceived by M.N.W and D.K. M.N.W, E.K., M.T., E.Z. and D.K. were responsible for study design. M.N.W. fabricated flexible 3D interfaces with contributions from M.R. and F.S. and performed multimodal experiments. S.H. performed finite element modeling. M.N.W., S.H., C.W.C., I.S., E.C. and D.K. performed interface characterizations and analysis. E.K.K. generated cortical organoids. J.L. maintained organoid-interface co-cultures and contributed to multimodal experiments. E.K.K. and S.E. performed immunohistochemistry and quantifications. M.S. performed transcriptomic analysis. F.M. performed CLARI-O immunostaining. M.N.W., J.Y., J.L. and D.K. processed and analyzed multimodal experimental data. M.N.W., E.K.K., J.Y., J.L., M.R., T.Z., E.C., M.T., E.Z. and D.K. contributed to data analysis and interpretation. The paper was written by M.N.W. and D.K. with input from E.K.K., E.C., M.T. and E.Z. All authors approved the final manuscript.

## Competing Interests

The authors have no competing interests to declare.

## Data availability

The data supporting the findings of this study are available within the paper and its Supplementary Information. Source data are provided with this paper in the Source data file. Other relevant data are available upon request from the corresponding author.

## Code availability

The code used for processing the data is available from the corresponding author upon request.

## Notes

### Competing Interest Statement

The authors have declared no competing interest.

